# The deubiquitinase Usp9x regulates PRC2-mediated chromatin reprogramming during mouse development

**DOI:** 10.1101/2020.06.28.176412

**Authors:** Trisha A. Macrae, Miguel Ramalho-Santos

## Abstract

Pluripotent cells of the mammalian embryo undergo extensive chromatin rewiring to prepare for lineage commitment after implantation. Repressive H3K27me3, deposited by Polycomb Repressive Complex 2 (PRC2), is reallocated from large gene-distal blankets in pre-implantation embryos to mark promoters of developmental genes. The factors that mediate this global redistribution of H3K27me3 are unknown. Here we report a post-translational mechanism that destabilizes PRC2 to constrict H3K27me3 during lineage commitment. Using an auxin-inducible degron system, we show that the deubiquitinase Usp9x is required for mouse embryonic stem (ES) cell self-renewal. Usp9x-high ES cells have high PRC2 levels and bear a chromatin and transcriptional signature of the pre-implantation embryo, whereas Usp9x-low ES cells resemble the post-implantation, gastrulating epiblast. We show that Usp9x interacts with, deubiquitinates and stabilizes PRC2. Deletion of *Usp9x* in post-implantation embryos results in the derepression of genes that normally gain H3K27me3 after gastrulation, followed by the appearance of morphological abnormalities at E9.5, pointing to a recurrent link between Usp9x and PRC2 during development. Usp9x is a marker of “stemness” and is mutated in various neurological disorders and cancers. Our results unveil a Usp9x-PRC2 regulatory axis that is critical at peri-implantation and may be redeployed in other stem cell fate transitions and disease states.

## INTRODUCTION

Immediately after implantation, the pluripotent embryonic epiblast enters a period of accelerated growth. This amplification event corresponds to a transition in cell fate from a pre-implantation state of naïve pluripotency to a post-implantation state of lineage priming. The stages of pluripotency can be modeled in vitro using mouse Embryonic Stem (ES) cells: culture in dual Mek/Gsk3β inhibition with leukemia inhibitory factor (LIF) and vitamin C maintains a pre-implantation-like state of pluripotency^1–3^, while ES cells in serum/LIF mimic the fast-growing state of early post-implantation epiblast cells^4,5^.

Reprogramming of the chromatin landscape contributes to the transition in pluripotent cell states at peri-implantation^6^. This reprogramming event includes a global redistribution of the repressive histone mark H3K27me3, deposited by Polycomb Repressive Complex 2 (PRC2). Recent studies document that H3K27me3 marks broad genic and intergenic domains in pre-implantation embryos as well as naïve ES cells^7–9^. After implantation, H3K27me3 becomes concentrated over promoters of developmental regulatory genes^7,10^, resembling patterns that restrain expression of bivalent (H3K27me3/H3K4me3-marked) genes in serum ES cells^11,12^. The mechanisms that regulate this peri-implantation switch in PRC2 activity are unknown.

We recently reported a genome-wide screen that revealed that the chromatin state of ES cells is acutely tuned to variations in protein synthesis and degradation^5^. The deubiquitinating enzyme Ubiquitin Specific Protease 9x (Usp9x) was one of the top hits in this screen. Although its roles in chromatin regulation have not been investigated, Usp9x is a marker of “stemness” ^13,14^ and is a key, conserved regulator of several stem/progenitor cells, including neural^15–18^, hematopoietic^19^, muscle^20^ and intestinal cells^21^. For example, Usp9x promotes self-renewal of mouse neural stem/progenitor cells^15,18^, and *USP9X* mutations are implicated in X-linked neurodevelopmental syndromes^22–24^, Turner Syndrome^25^, intellectual disability^26^ and seizures^27^ (reviewed in ref. 28). Moreover, *USP9X* mutations occur frequently in human cancers^29–31^. We report here that Usp9x deubiquitinates and stabilizes PRC2, acting as a gatekeeper to the switch in H3K27me3 deposition patterns during mouse development. These findings shed light on the regulation of chromatin reprogramming in pluripotent cells and during lineage commitment, and have important implications for physiological and pathological settings where Usp9x and PRC2 have been shown to play roles.

## RESULTS

### Usp9x promotes ES cell self-renewal and a transcriptional state of pre-implantation

We established an auxin inducible degron (AID) system for acute control of Usp9x protein levels^32,33^ (Fig. 1a). In ES cells homozygous for the *OsTir1* auxin receptor, we tagged endogenous *Usp9x* with enhanced green fluorescent protein (GFP) and a minimal AID or 3x Flag tag (herein referred to as AID-Usp9x or Flag-Usp9x, respectively). Auxin drives substantial Usp9x protein depletion in AID-Usp9x cells within approximately 8 hours (h) (Supplementary Fig. 1a). We used GFP expression to isolate subpopulations that resist degradation (Usp9x-high) or lose Usp9x (Usp9x-low) in response to auxin (Fig. 1a), each fraction corresponding to ∼20% of the total population.

**Figure 1.**
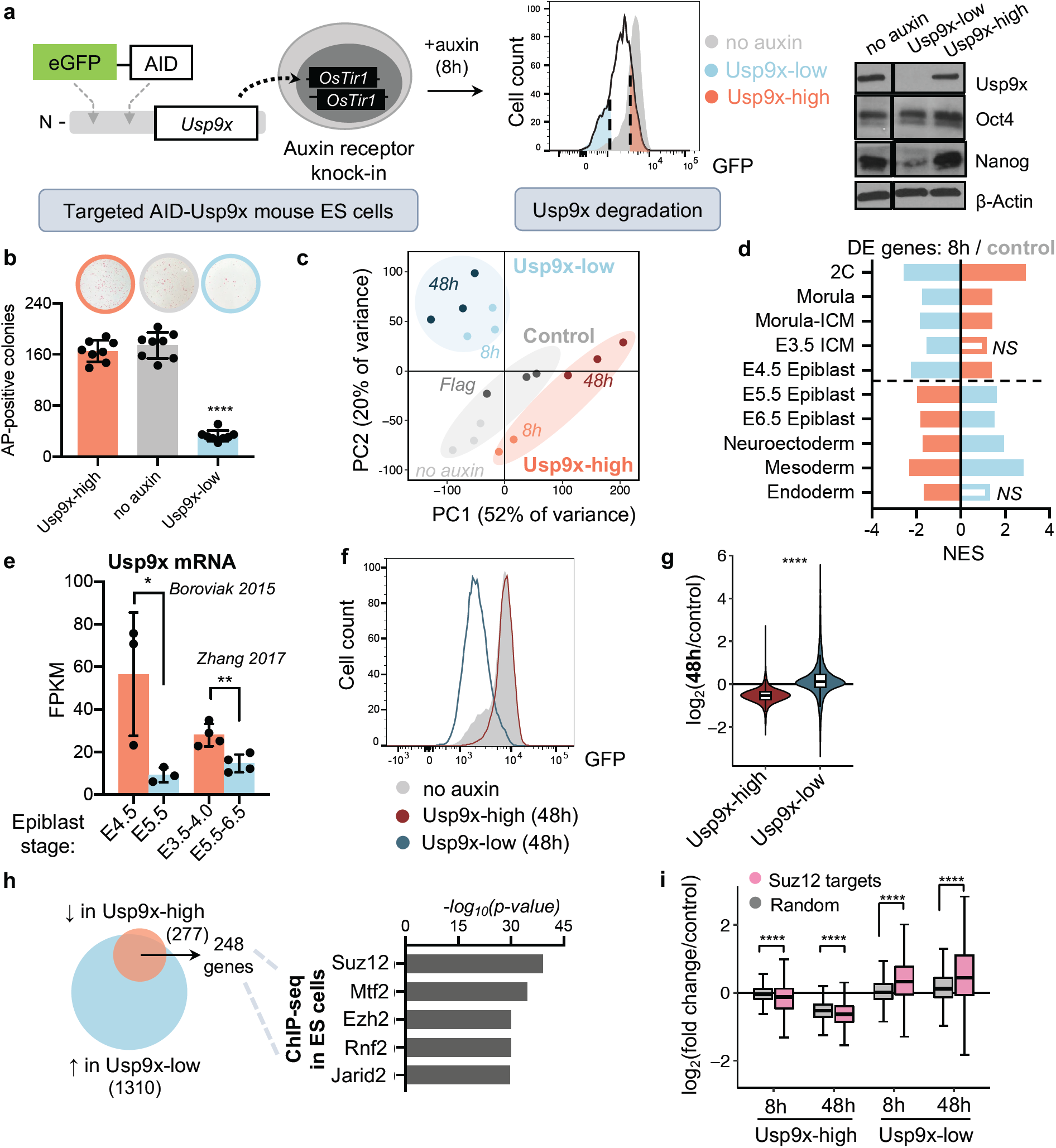
Usp9x promotes ES cell self-renewal and a transcriptional signature of pre-implantation linked to PRC2 activity. **a)** Schematic of an auxin-inducible degron (AID) system for acute Usp9x depletion in mouse embryonic stem (ES) cells. Right: western blot confirming endogenous Usp9x depletion in Usp9x-low and retention in Usp9x-high ES cells. **b)** Usp9x-low ES cells show a self-renewal deficit. Representative images and quantification of colony formation assays. *AP*, Alkaline Phosphatase. **c)** Principal Component Analysis (PCA) of gene expression by RNA-seq. *8h*: 8h auxin. *No auxin*: AID-Usp9x cells with vehicle treatment. *48h*: 8h auxin followed by 48h recovery without auxin. *Flag*: Flag-Usp9x cells after 8h auxin and 48h recovery. **d)** The transcriptional signatures of Usp9x-high or Usp9x-low ES cells correlate with different stages of peri-implantation development by Gene Set Enrichment Analysis (GSEA). See Methods for references. *NES*, Normalized Enrichment Score, *DE*, differentially expressed; *NS*, not significant (FDR > 0.05). **e)** Usp9x mRNA expression in the epiblast declines from pre- to post-implantation^36,37^. **f)** Usp9x-low ES cells do not recover Usp9x expression after 48h without auxin. **g)** Violin plots of the fold-change in expression of all genes at 48h relative to control cells, showing hypotranscription in Usp9x-high ES cells and hypertranscription in Usp9x-low ES cells. **h)** The overlap of genes DE in Usp9x-high and Usp9x-low ES cells are enriched for PRC2 binding by Enrichr analysis^42^. **i)** Boxplots showing repression (in Usp9x-high) or induction (in Usp9x-low) of Suz12 target genes^49^, compared to a random subset (*n* = 3350). Data are mean ± s.d. of 4 replicates from 2 sorts (b), mean ± s.d. (e), mean ± s.e.m. (i). Boxplot hinges (h, i) show the first and third quartiles, with median center line. \**P* < 0.05, \*\**P* < 0.01, ****P < 2.2 x 10^−16^. *P*-values by one-way ANOVA with multiple t-test comparisons to the no-auxin condition (b), Student’s t-test with Welch’s correction (e), Wilcoxon rank-sum test (g), and ANOVA with multiple pairwise Wilcoxon tests (i).

Usp9x-high and Usp9x-low ES cells express comparable levels of Oct4, but Usp9x-low ES cells are Nanog-low and display a 5-fold reduction in self-renewal capacity (Fig. 1b). Knockdown of Usp9x by an alternative method (RNA interference) also induces loss of self-renewal (Supplementary Fig. 1b). Furthermore, Usp9x expression declines with early lineage commitment by Embryoid Body (EB) formation, and low Usp9x expression does not represent a distinct cell cycle stage (Supplementary Fig. 1c-e).

We performed cell number-normalized (CNN) RNA-sequencing (RNA-seq) with spike-ins to characterize Usp9x-high and Usp9x-low ES cells. By principal component analysis (PCA), replicates cluster according to Usp9x levels (Fig. 1c). We calculated differential expression in 8h Usp9x-high or Usp9x-low ES cells versus controls and compared their profiles to molecular signatures of development using Gene Set Enrichment Analysis (GSEA)^34^ (Supplementary Table 1). This analysis revealed a striking polarity based on Usp9x levels: the Usp9x-high state correlates with pre-implantation embryonic stages, whereas Usp9x-low ES cells resemble the post-implantation epiblast and early lineages (Fig. 1d). Usp9x-high ES cells express high levels of naïve state markers and low levels of primed state markers^5,35^, while the opposite is observed in Usp9x-low ES cells (Supplementary Fig. 1f, g). Moreover, the expression of Usp9x itself declines from pre- to post-implantation in wild-type embryos (Fig. 1e, Supplementary Fig. 1h)^36,37^. These results indicate that Usp9x promotes ES cell self-renewal and that loss of Usp9x captures the transcriptional reprogramming that occurs in pluripotent cells at implantation.

ES cells cultured in serum/LIF represent a heterogeneous mixture of interconvertible pluripotent states. Surprisingly, isolated Usp9x-low cells do not re-distribute along a spectrum of Usp9x expression after a 48h recovery period without auxin (Fig. 1f and Supplementary Fig. 2a), unlike the cases of naïve pluripotency markers such as Nanog or Rex1^38,39^. Usp9x-high cells settle into a state of *hypo*transcription^4^, demonstrating a suppression of the majority of the transcriptome relative to control cells. By contrast, Usp9x-low cells at 48h show relative *hyper*transcription as well as induction of differentiation- and development-related Gene Ontology (GO) terms (Fig. 1g and Supplementary Fig. 2b, c), similar to the predicted upregulation of transcriptional output and lineage induction in the post-implantation epiblast^5,40,41^.

We probed the Usp9x-associated transcriptional signatures for clues to the regulatory networks establishing such divergent cell fates. Consistent with their anti-correlated GSEA signatures (Fig. 1d), Usp9x-high and Usp9x-low ES cells show polarized expression of many of the same genes. Of the 277 genes significantly downregulated in Usp9x-high cells, 248 (90%) are significantly upregulated in Usp9x-low cells. ChIP-X Enrichment Analysis (ChEA) revealed that these genes are enriched for targets of Polycomb Repressive Complex 2 (PRC2) (Fig. 1h and Supplementary Fig. 2d)^42,43^. Deletion of core members (Suz12, Ezh2, or Eed) in ES cells leads to induction of developmental regulatory genes^44,45^ and can promote premature lineage commitment^46–48^, similar to the behavior of Usp9x-low ES cells (Supplementary Fig. 1f). These data indicate that Usp9x-high ES cells represent a PRC2-repressed, pre-implantation state of pluripotency, while activation of a subset of PRC2 targets in Usp9x-low ES cells promotes a post-implantation state of lineage induction (Fig. 1i).

### *Usp9x*-mutant embryos arrest at mid-gestation with incomplete repression of a subset of PRC2-targeted lineage genes

We next turned to a mouse model to study the role of Usp9x in developmental progression. *Usp9x*-mutant embryos arrest at mid-gestation^50,51^, although the stage and underlying causes of developmental arrest are unknown. To avoid confounding effects from roles of Usp9x in cleavage-stage and trophectoderm development^52,53^, we used a *Sox2-Cre* to delete *Usp9x* strictly in the post-implantation epiblast derivatives of embryos^54^. We then genotyped and catalogued the morphology of control (ctrl, *Usp9x*^*fl/Y*^) versus mutant (mut, *Usp9x*^*Δ/Y*^) embryos at several mid-gestation stages (Fig. 2a, Supplementary Fig. 3a). Deviation from the expected (1:1) ratio arises by E11.5, at which point mutants account for only ∼25% of recovered embryos and have morphological abnormalities with 100% penetrance. The few mutants that survive to E11.5 show extensive hemorrhaging, while most display pericardial edema, cerebral edema and severe delay, pointing to an earlier developmental arrest (Fig. 2b). *Usp9x* mutants already display developmental delay (delayed turning, open anterior neural tube) or gross abnormalities by E9.5, including blunted posterior trunk development and exencephaly (Fig. 2b). These pleiotropic outcomes agree with the phenotypes of E9.5 chimeric embryos derived from *Usp9x*-genetrapped ES cells and the ubiquitous expression of Usp9x at E9.5 ^51,55^.

**Figure 2.**
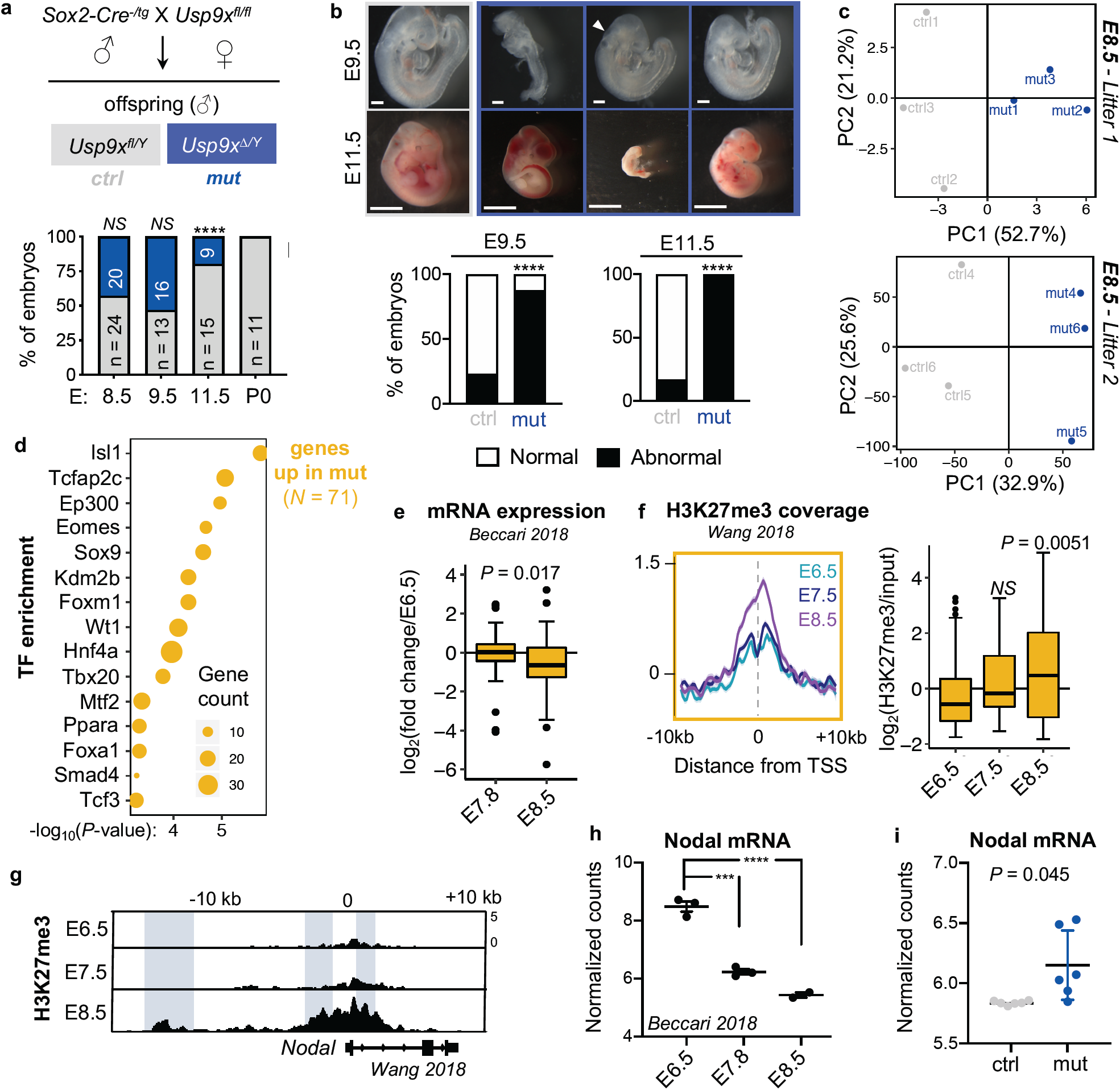
*Usp9x*-mutant embryos arrest at E9.5-E11.5 and display defective repression of early lineage programs marked by H3K27me3. **a)** Genetic cross to delete *Usp9x* in epiblast derivatives of post-implantation embryos. Below, quantification of male (XY) embryos at several post-implantation stages. **b)** Representative images and quantification of control and mutant embryo phenotypes. Scale bars = 250 µm (E9.5), 2.8 mm (E11.5). **c)** PCA plots of RNA-seq from litter-matched mutants and controls, showing that genotypes separate along PC1. **d)** Enrichr analysis of the top-enriched transcription factors (TF) that bind to the genes upregulated in *Usp9x*-mutant embryos in various cell types. **e)** Expression of the genes upregulated in *Usp9x* mutants during wild-type development^56^. **f)** Distribution and boxplot quantification of H3K27me3 levels over the genes upregulated in *Usp9x* mutants^58^. **g)** Representative genome browser tracks of H3K27me3 in wild-type embryos (E6.5-E8.5) at the *Nodal* locus^58^. Known enhancer elements are highlighted and show gains of H3K27me3. **h)** Nodal mRNA expression in wild-type development^56^. **i)** Nodal derepression in *Usp9x-*mutant versus control embryos at E8.5. Boxplot hinges (e, f) show the first and third quartiles, with median center line. Data are mean ± s.e.m. (h, i). \*\*\**P* < 0.001, \*\*\*\**P* < 0.0001 or indicated. *P*-values obtained by *χ*^2^ test (a, b), Student’s t-test with Welch’s correction (e, i), Wilcoxon rank-sum test (f), and ANOVA with multiple t-test comparison to E6.5 (h).

*Usp9x* mutants appear morphologically normal at E8.5 (Supplementary Fig. 3b). We therefore performed whole-embryo RNA-seq at this stage to identify early transcriptional changes that may anticipate subsequent developmental defects. As expected, the transcriptional differences are relatively minor at this stage: we identified 71 upregulated and 66 downregulated genes in *Usp9x* mutants (Supplementary Fig. 3c,d and Supplementary Table 2). Nevertheless, E8.5 *Usp9x* mutants are readily distinguished from controls by PCA and unsupervised hierarchical clustering (Fig. 2c, Supplementary Fig. 3e). Upregulated genes in *Usp9x* mutants are also upregulated in 48h Usp9x-low ES cells (Supplementary Fig. 3f). These genes include targets of master developmental transcription factors (Isl1, Tfap2c, Eomes, Sox9, Hnf4a, among others, Fig. 2d), and are enriched for processes in cardiac/mesoderm and endoderm development (Supplementary Fig. 3g). Overall, the genes upregulated in *Usp9x-*mutants at E8.5 typically decline by this point during wild-type development^56^ (Fig. 2e), suggesting that Usp9x is required for appropriate silencing of developmental regulatory genes.

Incomplete repression of regulators of lineage commitment with defective differentiation is also observed in PRC2-hypomorphic ES cells^49,57^. Recent chromatin immunoprecipitation-sequencing (ChIP-seq) of wild-type mouse embryos documented a wave of H3K27me3 deposition during gastrulation^58^. We therefore probed the developmental dynamics of H3K27me3 levels at genes differentially expressed in *Usp9x* mutants. Interestingly, the genes upregulated in E8.5 *Usp9x* mutants normally gain significant amounts of H3K27me3 between E6.5 and E8.5 (Fig. 2f), suggesting that PRC2 activity contributes to repressing them (Fig. 2e) at this stage. The TGFβ superfamily member *Nodal* is one key gene that normally accumulates H3K27me3 concurrent with its downregulation by E8.5 (Fig. 2g,h). Nodal remains upregulated in E8.5 *Usp9x* mutants (Fig. 2i). We speculate that persistent expression of earlier developmental regulators such as Nodal may impede developmental progression of *Usp9x* mutant embryos. Consistent with this notion, the 66 genes *down*regulated in mutants are normally induced from E7.5-E8.5 (Supplementary Fig. 3h-j). Taken together, these results indicate that E8.5 *Usp9x* mutant embryos display incomplete repression of early post-gastrulation lineage commitment genes that normally gain H3K27me3 at this stage.

### Usp9x mediates a pre- to post-implantation switch in H3K27me3 distribution

Our transcriptional analyses point to a role for Usp9x in promoting PRC2-mediated silencing of developmental regulatory genes, both in ES cells to prevent their premature activation prior to lineage commitment (Fig. 1) and in post-gastrulation embryos to allow for subsequent development (Fig. 2). We returned to ES cells to dissect the mechanism by which Usp9x regulates PRC2 activity. Consistent with their signature of PRC2 target gene derepression (Fig. 1i), Usp9x-low ES cells display globally reduced levels of H3K27me3 compared to Usp9x-high ES cells by cell number-normalized western blot (Fig. 3a). We next used spike-in normalized ChIP-seq to map the genome-wide levels and distribution of H3K27me3 between Usp9x-associated cell states. Compared to Usp9x-low ES cells, Usp9x-high ES cells display global gains of H3K27me3 at bivalent (dual H3K4me3/H3K27me3-marked) promoters^59^ (Supplementary Fig. 4a-c), which are canonical PRC2 targets highly enriched for developmental regulators. H3K27me3 gains are not limited to bivalent promoters, as Usp9x-high cells carry higher levels over peaks present at baseline (no-auxin condition) and spreading upstream and downstream of peaks (Fig. 3 b,c and Supplementary Fig. 4d). Usp9x-high cells also have H3K27me3 enrichment over repetitive elements (Fig. 3d), which are targeted by this mark in naïve, pre-implantation-like conditions^60^. Thus, the transition from Usp9x-high to Usp9x-low ES cells involves a genome-wide reduction in H3K27me3 and a narrowing of its peaks across developmental genes and repeat elements.

**Figure 3.**
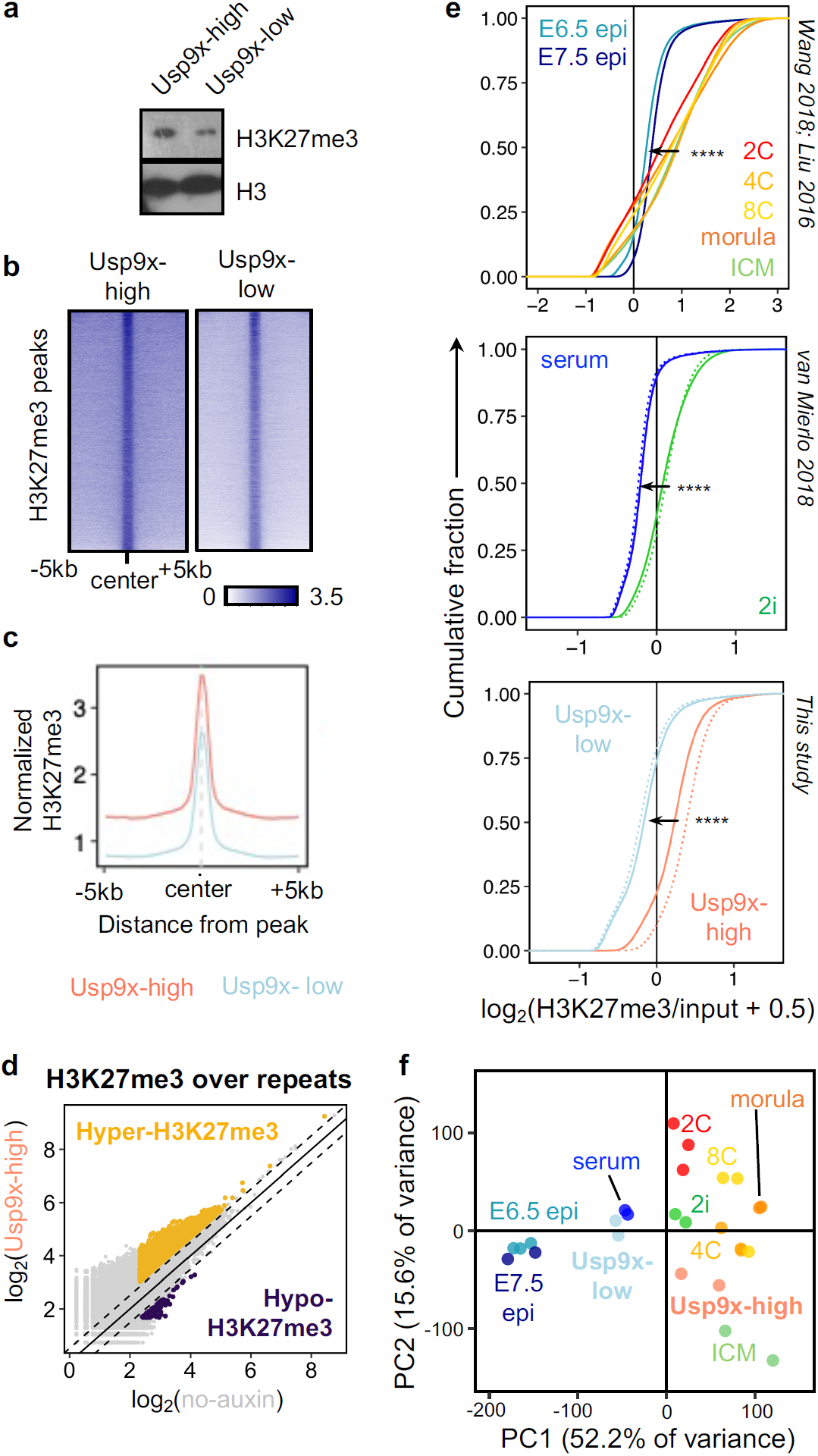
Usp9x mediates a pre- to post-implantation switch in H3K27me3 distribution. **a)** CNN western blot of H3K27me3 from histone extracts, representative of 2 biological replicates. **b)** Heatmaps of H3K27me3 ChIP-seq signal in Usp9x-high and Usp9x-low ES cells, showing H3K27me3 spreading in Usp9x-high cells. **c)** Profile plot depicting the mean signal of coverage shown in (b). **d)** Usp9x-high ES cells carry more H3K27me3 over repetitive elements compared to untreated (no-auxin) cells. Each point represents an individual element. **e)** Cumulative enrichment plots of H3K27me3 enrichment in non-overlapping genomic bins of *in vivo* developmental stages (top) and ES cell states (middle, bottom)^8,58,65^. Pre-implantation stages (2C-ICM, top) or pre-implantation-like ES cell states (2i, Usp9x-high) display H3K27me3 enrichment. *Epi*, epiblast. **f)** PCA plots clustering Usp9x-high and Usp9x-low ES cells among the samples shown in (e) based on H3K27me3 distributions. Each point represents a biological replicate. **** *P* < 2.2 x 10^−16^ by Kolmogorov-Smirnov test.

The pattern of H3K27me3 in Usp9x-high serum ES cells resembles the diffuse domains of the mark in naïve ES cells under dual Mek/Gsk3β inhibition (2i) and in pre-implantation embryos^7–9,61^. In agreement with this notion, cumulative enrichment plots revealed that the global drop in H3K27me3 levels in Usp9x-high versus Usp9x-low ES cells recapitulates what is observed in the transition from 2i to serum ES cells and from pre-implantation to post-implantation embryos (Fig. 3e). PCA of H3K27me3 ChIP-seq data separates pre-implantation and post-implantation embryos along PC1. ES cell data also follow this trajectory, with Usp9x-high and 2i ES cells aligning with pre-implantation and Usp9x-low and serum ES cells clustering with post-implantation stages (Fig. 3f). These results indicate that H3K27me3 enrichment across large swaths of the genome (Fig. 3), together with a PRC2-repressed transcriptional program (Fig. 1), are hallmarks of pre-implantation pluripotency^62–64^ and Usp9x-high ES cells.

### Usp9x deubiquitinates and stabilizes PRC2

We next explored the possibility that Usp9x may directly regulate PRC2 levels or activity. We found that the protein levels of core PRC2 components are downregulated in Usp9x-low ES cells (Fig. 4a). This finding led us to hypothesize that Usp9x deubiquitinates and stabilizes PRC2 components to drive H3K27me3 deposition. In support of this notion, endogenous Usp9x interacts with Suz12 and Ezh2 in ES cell nuclear extracts (Fig. 4b, Supplementary Fig. 5a). Moreover, acute AID-Usp9x depletion leads to the accumulation of poly-ubiquitinated forms of Suz12 and Ezh2, within 4-8h of auxin addition (Fig. 4c). Alternative methods of reducing Usp9x activity, either small molecule inhibition (WP1130)^66^ or overexpression of a mutant catalytic domain (C1566S), also lead to accumulation of ubiquitinated forms of Suz12 and Ezh2 (Fig. 4d and Supplementary Fig. 5b,c). These gains of ubiquitin upon Usp9x loss correlate with destabilization of Suz12 and/or Ezh2 (Fig. 4a,c-d and Supplementary Fig. 5b,c). Taken together, these data indicate that Usp9x interacts with PRC2 components and that its catalytic activity is required to promote a deubiquitinated state and higher protein levels of PRC2.

**Figure 4.**
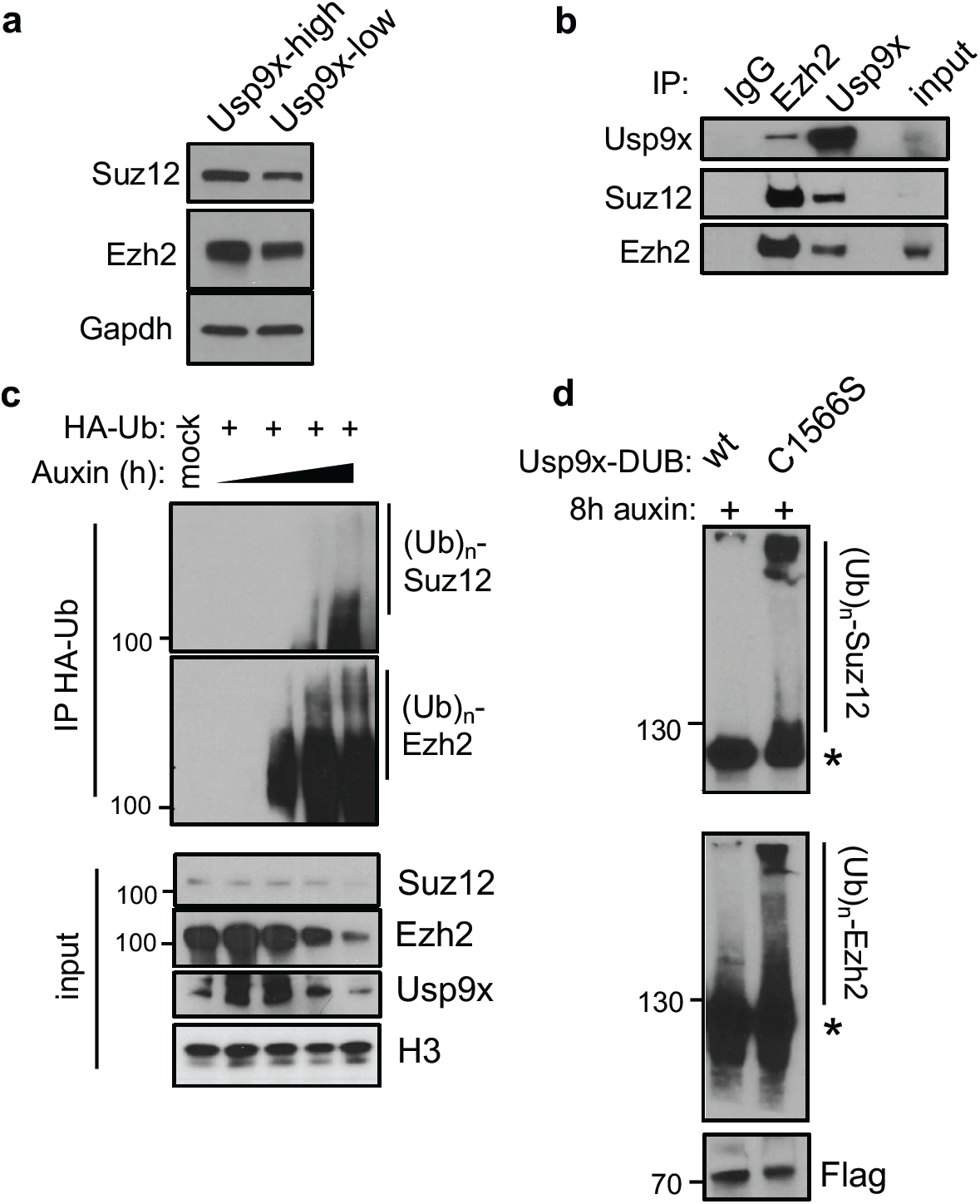
Usp9x is a PRC2 deubiquitinase. **a)** CNN western blots for Suz12 and Ezh2 proteins in whole cell extracts. **b)** Co-immunoprecipitation (*IP*) and western blot showing a reciprocal interaction between endogenous Ezh2 and Usp9x in wild-type ES cells. **c)** Acute auxin depletion over a time course from 0-8h leads to gain of ubiquitinated species of Suz12 and Ezh2 and destabilization of their protein levels. *HA-Ub*, HA-tagged ubiquitin; *(Ub)*_*n*_, polyubiquitination. **d)** Overexpressing a catalytic-mutant (C1566S) versus wild-type (wt) form of the Usp9x catalytic domain (DUB) leads to gain of Suz12 and Ezh2 ubiquitin levels. AID-Usp9x cells were treated with auxin for 8h to deplete endogenous Usp9x. Asterisk (*) designates the expected sizes for non-ubiquitinated species. Western blots are representative of at least 2 biological replicates.

## DISCUSSION

In summary, we report here that Usp9x deubiquitinates core PRC2 members to promote high levels of H3K27me3, repress developmental regulatory genes and maintain a pre-implantation-like state in ES cells (Fig. 5). Studies in mammalian systems have emphasized the developmental role of PRC2 in regulating bivalent promoters^11,12^, which represent the highest-affinity sites for PRC2 activity^61,67,68^. However, H3K27me3 is widespread outside of bivalent chromatin over the genomes of pre-implantation embryos and Usp9x-high ES cells, and broad domains occur in other cell types later in development^69–72^. Promiscuous activity may be an ancestral function of PRC2^73^. While the mechanisms underlying such promiscuity remain unclear, the Eed subunit has been shown to promote spreading of H3K27me3 domains^74^. PRC2 stability may also be a major factor^75^. Oncogenic *EZH2* mutations stabilize the complex and cause ectopic gains of H3K27me3 in lymphoma^76–78^. Our results highlight a mechanism whereby Usp9x-stabilizes the PRC2 complex to promote global increases in H3K27me3 and expansion to lower-affinity sites (Fig. 5). It will be of interest to determine how the partnership between Usp9x and PRC2 integrates with the activity of other Usp9x substrates, as well as to identify the E3 ubiquitin ligase(s) acting on PRC2 during development.

**Figure 5.**
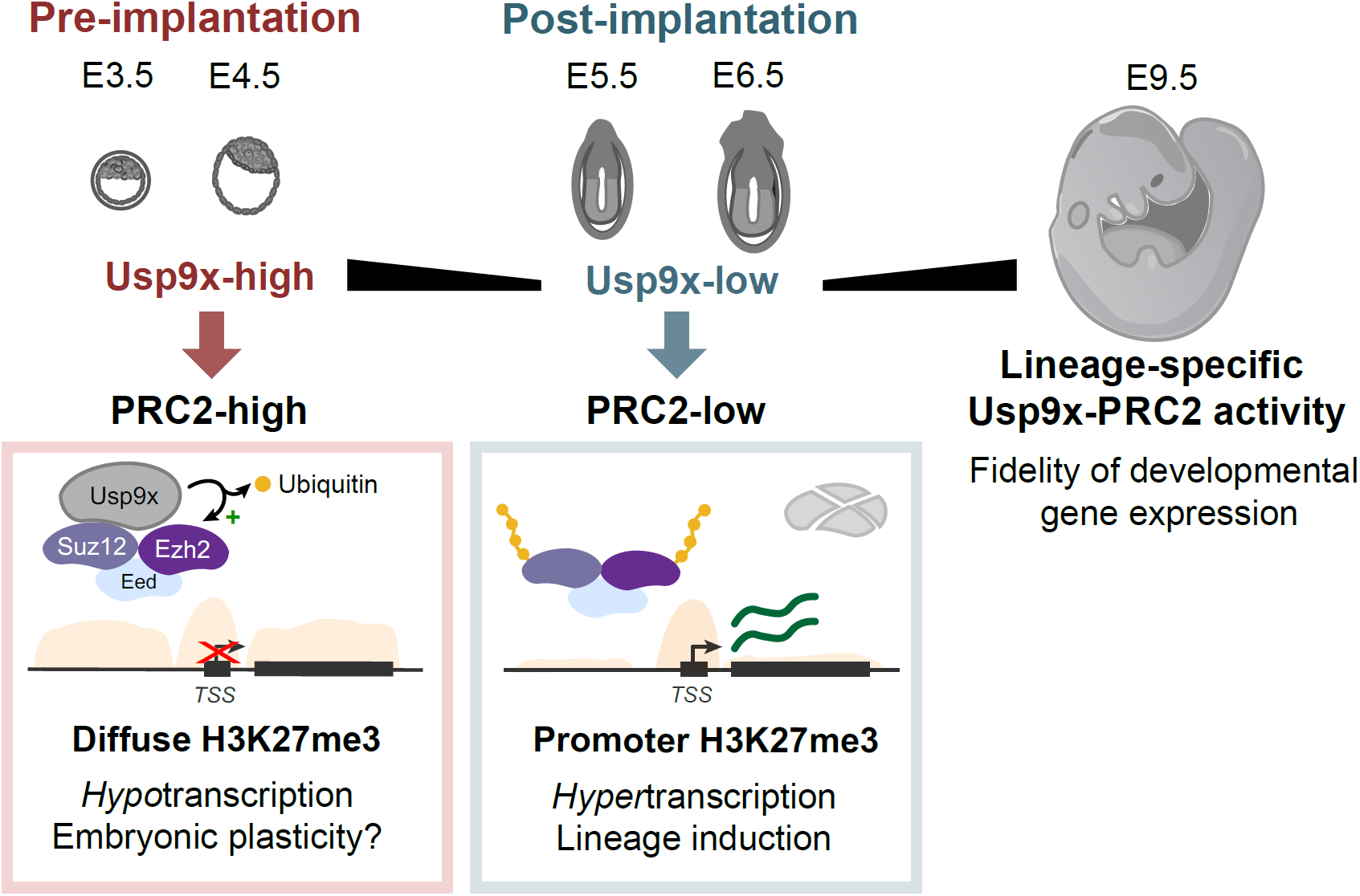
Model for the Usp9x-PRC2 regulatory axis in early development. Usp9x is a PRC2 deubiquitinase that promotes diffuse H3K27me3 deposition and a pre-implantation-like transcriptional state. Loss of Usp9x leads to PRC2 destabilization, restricted H3K27me3 deposition and global hypertranscription with priming of post-implantation lineages. After gastrulation, *Usp9x* is required for timely silencing of developmental genes that are PRC2 targets and for normal development beyond E8.5.

Maternally-inherited H3K27me3 domains have been proposed to mediate non-canonical genomic imprinting in mouse embryos^79,80^ and restrict enhancer expression in early-stage fly embryos^81^. Our finding that Usp9x-high/PRC2-high ES cells enter a state of global hypotranscription (Fig. 1f,i and Supplementary Fig. 2b) raises the possibility that ubiquitous H3K27me3 *in vivo* suppresses large-scale transcription prior to implantation (Fig. 5), possibly by preventing H3K27 acetylation^43,82,83^. Extensive H3K27me3 may also safeguard embryonic potential during segregation of extraembryonic lineages^62^.

Our data suggest that the decline in Usp9x expression at implantation contributes to destabilizing PRC2 to allow lineage induction. The roles of PRC2 after gastrulation remain obscure. In zebrafish and Xenopus, H3K27me3 marks spatially-regulated genes after gastrulation^84,85^. Constitutive *PRC2* knockouts in mouse are peri-gastrulation lethal^86–88^, but these findings are confounded by requirements for PRC2 in extraembryonic tissues^89^, and the developmental consequences of epiblast-targeted PRC2 knockouts are unknown. Intriguingly, recent studies indicate that re-establishment of H3K27me3 after gastrulation may also regulate spatial gene expression in mouse^90,91^. Together with the data presented here, these results suggest that Usp9x and PRC2 are re-deployed to promote timely H3K27me3 deposition and silencing of key developmental genes after lineage commitment (Fig. 5). Further studies are required to understand how Usp9x may regulate batteries of PRC2 target genes in a lineage-specific manner during organogenesis.

The transition from a pre-implantation to a post-implantation state of pluripotency mirrors stem cell expansion events in other compartments. Supporting the designation of Usp9x as a “stemness” factor^13,14^, loss-of-function studies document that Usp9x restricts premature expansion of embryonic (this study), neural^17,18^, hematopoietic^19^ and intestinal stem/progenitor cell compartments^21^ in mice. Similarly, PRC2 plays essential roles in each of these stem cell compartments^92–95^. Mutations in *USP9X* are associated with several neurodevelopmental and neurodegenerative disorders as well as cancers ^22,23,26–31,96^. Thus, the Usp9x-PRC2 axis reported here merits further exploration in other developmental contexts and disease states.

## ACKNOWLEDGMENTS

We thank Aydan Bulut-Karslioglu and members of the Santos laboratory for input and critical reading of the manuscript. We are grateful to Elphège Nora and Benoit Bruneau for the AID targeting vector and *OsTir1*-knockin ES cells. We also thank J. Lee and the Toronto Center for Phenogenomics for mouse colony maintenance and support; M. Abed and V. Dixit for sharing the *Usp9x*-flox mice; E. Chow, K. Chan, and members of the UCSF Center for Advanced Technology and the LTRI Sequencing Core for next-generation sequencing; D. Gökbuget for sonication assistance; S. Biechele for cloning advice; and M. Percharde for bioinformatics guidance. Flow cytometry and sorting were performed in the UCSF Parnassus Flow Cytometry Core, supported by a Diabetes Research Center Grant and National Institutes of Health (NIH) grant P30 DK063720. This work was supported by NIH grant 1F30HD093116 (to T.A.M.) and NIH grants R01GM113014 and R01GM123556, a Canada 150 Research Chair in Developmental Epigenetics, and the Medicine by Design Program of the University of Toronto (to M.R.-S).

## AUTHOR CONTRIBUTIONS

T.A.M. and M.R.-S. conceived of the project. T.A.M. designed, performed, and analyzed all experiments. M.R.-S. supervised the project. T.A.M. and M.R.-S. wrote the manuscript.

## CODE AVAILABILITY

Custom R codes used for data analysis are available upon request.

## DATA AVAILABILITY

RNA-seq and ChIP-seq data have been deposited in Gene Expression Omnibus (GEO).

## METHODS

### Mice

*Usp9x*^*fl/fl*^ females were maintained as homozygotes on a C57BL/6 background by crossing *Usp9x*^*fl/fl*^ and *Usp9x*^*fl/Y*^ mice^19^. Heterozygous male *Sox2-Cre* mice were obtained from Jackson Laboratories (JAX stock #008454) and bred with Cre-negative females to maintain a stock of heterozygous males ^54^. All mice were maintained on 12h light/dark cycle and provided with food and water *ad libitum* in individually ventilated units (Techniplast) in specific pathogen-free facilities at The Center for Phenogenomics, Toronto. All procedures involving animals were performed according to the Animals for Research Act of Ontario and the Guidelines of the Canadian Council on Animal Care. Animal Care Committee reviewed and approved all procedures conducted on animals at TCP (Protocol 22-0331). Sample size choice was not pre-determined.

Yolk sacs were dissected from embryos and used for DNA extraction with the Red Extract-N-Amp kit (Sigma). Usp9x status was assessed by PCR using Phire Green Hot Start II PCR Master Mix (Thermo Fisher Scientific). Cycling conditions: 98°C for 30 sec; 35 cycles of 98°C for 5 sec, 58°C for 5 sec, 72°C for 8 sec; 72°C for 1 min. See Supplementary Table 3 for primer sequences.

### Plasmid construction

An sgRNA was designed to target the *Usp9x* ATG with 20 nucleotide overhang in both directions. Cloning was performed by annealing pairs of oligos into pSpCas9(BB)-2A-GFP (PX458) (modified from GFP to BFP by site directed mutagenesis), a gift from Feng Zhang (Addgene plasmid # 48138; http://n2t.net/addgene:48138; RRID:Addgene_48138)^97^. Plasmid identity was verified by Sanger sequencing.

The eGFP-AID-Usp9x plasmid was assembled from a pEN458-eGFP-AID[71-114]-eGFP-CTCF N-terminal targeting construct (a gift from the B. Bruneau lab). The N-terminus was chosen for targeting due to repetitive sequences at the *Usp9x* C-terminus. The vector was digested with NruI, NsiI and XhoI and gel purified to remove regions of *CTCF* homology (Qiagen Gel Extraction kit). ∼900 bp homology arms to *Usp9x* were amplified from mouse genomic DNA using PrimeStar GXL polymerase (Takara, CA, USA) and Gibson assembly primers with 21 nucleotide overlap to adjacent fragments. The vector fragments and homology arms were cloned by Gibson Assembly (NEB HiFi Assembly Kit).

Oligos containing a 3xFLAG sequence were annealed by incubation in standard annealing buffer (10 mM Tris-HCl pH 7.5, 1 mM EDTA, 50 mM NaCl) for 5 min at 95°C followed by slow cooling to 25°C. The annealed fragment was then digested with BamHI and XhoI and cleaned up by MinElute PCR purification (Qiagen). The eGFP-AID-Usp9x vector was also digested with BamHI/XhoI and cleaned up by gel extraction (Qiagen). The 3xFlag sequence was then ligated into the digested eGFP-Usp9x plasmid (Takara DNA ligation kit #6023).

### ES cell targeting

Vectors were amplified by transformation into Stbl3 competent cells (Invitrogen). Resultant colonies were picked for miniprep DNA extraction (Qiagen) and screening by restriction enzyme digest. Positive clones were sequence-verified, purified by Maxiprep column extraction (Qiagen), concentrated by standard ethanol precipitation overnight, and used for nucleofection into low passage (p3) *OsTir1*-knockin ES cells derived as described^33^. Briefly, this line (148.4) was derived from E14 mouse ES cells and is homozygous for a Tir1-2A-Puro cassette (Addgene plasmid # 92142; http://n2t.net/addgene:92142; RRID:Addgene_92142) at the *TIGRE* locus.

5 million OsTir1 cells (passage 4) were nucleofected with 2.5 µg of the sgRNA plasmid and 20 µg of either eGFP-AID-Usp9x plasmid or eGFP-3xFLAG-Usp9x plasmid, using an Amaxa Nucleofector 2b device and ES nucleofection kit (Lonza) per the manufacturer’s instructions. Cells were diluted in 500 µl of medium and immediately plated onto 10 cm dishes with 10 ml pre-warmed medium. After 2 days, GFP-single and BFP-/GFP-double-positive cells were sorted by FACS and plated at clonal density (10,000 cells per 10cm dish). Clones were left to expand for 5 days before manual picking onto 96-well plates. Single clones were then dissociated and expanded for 2 days. Clones were screened for auxin responsiveness by replica plating onto 2 96w plates, addition of auxin to 1 plate, and measurement of eGFP fluorescence intensity by flow cytometry. Auxin-responsive clones were subsequently expanded and used as biological replicates for all analyses. Cells were periodically pulsed with puromycin (1 µg/ml for 1-2 days) to select against transgene silencing.

### Usp9x-CD-mCherry cloning

The Usp9x catalytic domain was amplified from a plasmid containing the full-length Usp9x ORF, obtained from DNASU ^98^; mCherry was amplified by PCR from a pcDNA3-mCherry plasmid. We then cloned the purified Usp9x-DUB and mCherry fragments into pEF1a-IRES-Neo, a gift from Thomas Zwaka (Addgene plasmid # 28019; http://n2t.net/addgene:28019; RRID:Addgene_28019), by Gibson Assembly. To make the C1566S catalytic mutant form of the Usp9x DUB domain, we performed site directed mutagenesis^99^. PCR was carried out with Phusion polymerase (New England Biolabs), with PCR cycling conditions: 98°C for 7 min; 12 cycles of 98°C for 30s, 61°C for 30s, 72°C for 3 min 45s; 3 cycles of 98°C for 30s, 56°C for 30s, 72°C for 3m 45s; 72°C for 10 min; 4°C hold. The PCR product was digested with DpnI for 3h at 37°C (New England Biolabs) and then 5 µl was transformed into Stbl3 competent cells (Thermo Fisher Scientific). See Supplementary Table 3 for primer sequences.

### Mouse embryonic stem cell culture

ES cells were passaged every 1-2 days and grown in standard ES-FBS (serum/LIF) medium: DMEM GlutaMAX with Na Pyruvate, 15% fetal bovine serum (Atlanta Biologicals, GA, USA), 0.1 mM Non-essential amino acids, 50 U/ml Penicillin/Streptomycin, 0.1 mM EmbryoMax 2-Mercaptoethanol and 1000 U/ml ESGRO supplement. Cells tested negative for Mycoplasma contamination.

Indole-3-acetic acid sodium salt (Sigma I5148-2G) was dissolved in water to 500 mM, filter sterilized, and stored as aliquots at −20°C. Stock solutions were thawed and diluted to 500 µM for all depletion experiments. Wild-type ES cells were used to determine the range of GFP-negative expression for sorting Usp9x-low ES cells upon auxin treatment. Usp9x-low and Usp9x-high cells correspond to the bottom and top ∼15-20% of the population by GFP expression, respectively.

### Embryonic stem cell differentiation

LIF was withdrawn from ES-FBS medium for spontaneous differentiation into Embryoid Bodies (EB). ES cells were counted, plated on low-attachment 6w plates, and harvested by trypsinization at day 2 and day 5 for qRT-PCR or western blot analysis.

### siRNA-mediated Knockdown

siRNA transfections were performed in ES cells using Lipfectamine 2000 and OptiMEM (Thermo Fisher Scientific). ES cells were plated 5-7h before transfection at a density of 5 x 10^5^ cells per well of a 6-well plate and transfected with 100 pmol siRNA, according to the manufacturer’s standard recommendations. A SMARTpool of 4 independent siRNAs were used to knock down Usp9x (Dharmacon), and a non-targeting siRNA (siGenome siControl #2, Dharmacon) was used as a control. Transfections were performed in ES-FBS medium without antibiotics, and the medium was replaced the next morning with complete ES-FBS. Cells were harvested for counting and colony formation assays or western blots 48h after transfection.

### Colony formation assay

1000 cells from the indicated conditions were sorted and plated onto a 12-well plate. 4 replicates were performed for 2 independent sorts. Cells were grown in self-renewal conditions (serum/LIF) for 5-6 days. Colonies were then washed 1x in PBS, fixed for 15 minutes at RT in 2% PFA, and stained according to the instructions of the VectorRed Alkaline Phosphatase (AP) Substrate Kit (Vector Laboratories, CA, USA). Colonies were manually scored based on colony morphology and AP staining (positive if >50% of colony area).

### qRT-PCR

cDNA synthesis was performed with the High-Capacity cDNA Reverse Transcription Kit (Thermo Fisher Scientific), using random hexamer priming for 2 hours at 37°C. KAPA 2x SYBR Green Master Mix, low ROX (KAPA) was used for qPCR and data were acquired on a QuantStudio 5 (Thermo Fisher Scientific) and analyzed in Prism v8 (GraphPad).

### RNA-seq library preparation

3 independent clones of each cell line (AID-Usp9x or Flag-Usp9x) were used for RNA-sequencing. Cells were plated the day before sorting, and auxin was added to a final concentration of 500 µM in fresh media for 8h. Cells were collected by trypsinization and resuspended in FACS buffer (10% FBS, PBS, ± 500 µM auxin) with SYTOX Blue (Thermo Fisher Scientific) for sorting. 250,000 cells from each condition were sorted on the basis of negative SYTOX Blue incorporation. For the 8h timepoint, sorted cells were immediately pelleted, resuspended in Buffer RLT + β-mercaptoethanol (Qiagen), snap frozen on dry ice, and stored at −80°C before library preparation. For the 48h recovery timepoint, cells were re-plated in regular ES-FBS medium, cultured for 48h in serum/LIF without auxin, lysed, and snap frozen. Sorts were performed on a Sony SH800 Single Cell Sorter (Sony).

RNA extractions from frozen lysates were performed on the same day using RNeasy Mini columns (Qiagen). Recovered total RNA was quantified by Qubit and quality was assessed using an Agilent Bioanalyzer, RNA pico kit (Agilent). Synthetic RNAs from the External RNAs Control Consortium (ERCC) Spike-in Mix1 (Thermo Fisher Scientific) were added at known concentrations to the same volume of RNA from the previous step, per manufacturer’s instructions (2 µl of 1:100 ERCC dilution added to 10 µl of RNA, representing ∼1-1.5 µg RNA). 1 µg of total RNA was used for mRNA isolation and library preparation using the NEBNext Ultra II Directional Library Prep Kit for Illumina with the mRNA Magnetic Isolation Module, per manufacturer’s instructions (New England Biolabs, NEB #E7420S and #E7490S). Library quality was assessed by Bioanalyzer High Sensitivity DNA chip (Agilent). Libraries were quantified by Qubit and pooled at equimolar concentration. Sequencing was performed on a HiSeq 4000 (Illumina) with 50 bp single-end reads at the UCSF Center for Advanced Technology.

For embryo RNA-seq, whole E8.5 embryos were dissected, cleaned of extraembryonic tissue, resuspended in buffer RLT + β-mercaptoethanol (Qiagen), and snap-frozen on dry ice. 3 litter-matched control and mutant embryos were collected from 2 litters, for a total of n = 12 individual embryos. RNA was extracted as above and 300 ng of total RNA was used for library preparation using the NEBNext Ultra II Directional Library Prep Kit for Illumina with the mRNA isolation module and NEBNext Multiplex Oligos for Illumina (New England Biolabs). DNA quality was assessed by Fragment Analyzer NGS (Agilent). Libraries were quantified by Qubit and pooled at equimolar concentration for sequencing on a NextSeq 500 (Illumina) with 75 bp single-end reads at the LTRI Sequencing Core.

### RNA-seq analysis

Libraries were trimmed of Illumina adaptor sequences and quality-checked using Trim Galore! (Babraham Bioinformatics), and then aligned to the mm10 transcriptome with ERCC sequences using TopHat2 v2.0.13 options -g 20 --no-coverage-search --library-type fr-firststrand --no-novel-indels^100^. Gene counts were obtained from featureCounts on the command line with options: -t exon -T 8 -s 2 -g gene_id. The table of raw counts was imported into R, filtered to remove low-count genes (genes with 0 counts in any sample and those with ≤ 3 counts per million, CPM by edgeR, across all samples were filtered out), and separated into ERCC and gene counts. Values for spike-in normalization were determined from ERCC counts corrected for overall library size using edgeR calcNormFactors (nf <-calcNormFactors(raw_ercc_counts, lib.size=N), where N <-colSums(raw_gene_counts). The CNN factors were then used to adjust gene counts using the limma-voom transformation (option lib.size = N*nf)^101^. Data were further analyzed and plotted using ggplot2. The threshold for significant differential expression was adjusted *P* < 0.05 and log2FC > |0.7| relative to control cells (AID-Usp9x without auxin and Flag-Usp9x). Boxplots and violin plots show fold-change relative to control cells obtained from toptable analyses.

For embryo RNA-seq, gene counts were obtained in the same manner, imported into R, and converted to a DESeq2 object (DESeqDataSetFromMatrix using sample information) for processing, DESeq2 version 1.24.0^102^. Genes with fewer than 10 counts across all samples were filtered out before differential expression analysis. Counts normalized by the DESeq2 rlog transformation were used for PCA and heatmaps of gene expression. Raw counts were used for differential expression analysis using the default parameters of the DESeq function. To account for staging differences between litters, we called differential expression between litter-matched mutants and controls, applied a statistical cutoff (adjusted *P* < 0.1), and overlapped the gene lists to obtain refined sets of up- or down-regulated genes. RNA-seq data from post-occipital embryos stages E6.5, E7.8, and E8.5 are from^56^. Published DESeq2 results were used to plot fold-changes in expression from E6.5, and raw fastq files were downloaded from NCBI GEO, converted to normalized gene counts as above, and used to plot Nodal expression.

### Gene Ontology analyses

Pathway analysis was performed by Gene Ontology (GO) analysis using DAVID 6.8 and geneontology.org^103–106^. Transcription factor binding enrichment of gene lists was performed with ChEA, part of the Enrichr suite (https://amp.pharm.mssm.edu/Enrichr/)^42,107^. Tables of enriched factors and P-values were downloaded and plotted in R.

Gene Set Enrichment Analysis (GSEA v6.0.12) was performed using the online GSEAPreranked tool (https://cloud.genepattern.org/gp/pages/login.jsf) with default conditions to compare differential expression (all genes sorted by log2FC) with gene sets from published datasets, outlined below^34^. Normalized enrichment scores were plotted in Prism v8 (GraphPad).

#### Datasets used for GSEA

The 2-cell embryo signature is from^108^. Transcriptional signatures from cleavage stages through E5.5 were retrieved from^36^, taking either the full gene list or the top 500 genes enriched for a particular time point from the published stage-specific expression analysis. The E6.5 epiblast signature was defined as genes upregulated in E6.5 epiblast relative to visceral endoderm and endoderm at E6.5^37^. A signature of early mesoderm was determined from published RNA-seq of ES cell differentiation, and we selected genes by fold-change of expression at the mesoderm stage compared to ES cells^109^. The endoderm signature comes from published microarray data of early endoderm in E7.5 embryos^110^. Neuroectoderm genes were defined by RNA-seq data of epiblast stem cell differentiation to neural fate, comparing the fold-change in expression at day 2 of differentiation relative to baseline^111^. In all cases, either the full published gene list or the top 500 genes ranked by fold-change were used for GSEA.

### Histone extraction

Histone extraction was performed using a standard acid extraction protocol^112^. Sorted cells were lysed for 10 min at 4°C in triton extraction buffer (PBS with 0.5% Triton X-100, 2 mM PMSF, 1x Halt Protease Inhibitor at a density of 10^7^ cells/ml). Lysates were spun for 10 min at 4°C, 2000 rpm. The pellet was washed once in 0.5x volume of lysis buffer and centrifuged again. Pellets were resuspended in 0.2 N HCl (10^6^ cells/ml) and acid extracted overnight, rotating at 4°C. The next day, the solution was clarified by centrifugation and the supernatant transferred to a new tube. Histones were precipitated in 0.25x volume TCA, incubated 20 minutes on ice, and pelleted at max speed for 10 min. Excess acid was removed from solution through two washes in ice-cold acetone, pellets were air-dried, and histones were resuspended in water for BCA Protein quantification (Pierce). LDS sample buffer (Thermo Fisher Scientific) was added to 1x and samples were denatured for 5 min at 95°C followed by cold shock.

### Co-immunoprecipitations

Co-immunoprecipitation (Co-IP) assays were performed on nuclear extracts. ES cells grown to ∼70% confluency were washed twice and then scraped in cold PBS. Cell pellets were weighed and resuspended in 4x volume of swelling buffer A (10 mM HEPES pH 7.9, 5 mM MgCl_2_, 0.25 M Sucrose, 0.1% NP-40) with protease inhibitors were added fresh (1x Halt Protease inhibitors, 1 mM PMSF, 1 mM NaF, 10 mM N-ethylmaleimide). Lysates were incubated on ice for 20 min and passed through a 18 1/2 G needle five times. Nuclei were pelleted by centrifuged for 10 min at 1500 *g* and lysed in 8x volume buffer B (10 mM HEPES pH 7.9, 1 mM MgCl_2_, 0.1 mM EDTA, 25% glycerol, 0.5% Triton X-100, 0.5 M NaCl with PIs as in buffer A). After incubation on ice for 10 min, samples were passed through an 18 ½ G needle 5 times and pulse sonicated using a probe sonicator, 2 times 5 seconds at 4°C. 100 µl of lysate was diluted in 400 µl IP wash/dilution buffer (150 mM NaCl, 10 mM Tris pH 8, 0.5% sodium deoxycholate, 1% Triton X-100, 1 mM EDTA, 1 mM EGTA) and rotated 4h-overnight with 1 µg Rb anti-Usp9x (Bethyl), 1.7 µg Rb anti-Ezh2 (CST #5246), or 1.7 µg Rb anti-IgG (Millipore CS200581). Input samples were collected at this time. Immune complexes were bound by 25 µl of pre-washed Protein A Dynabeads (Thermo Fisher Scientific), rotating end-over-end for 2h at 4°C. Beads were washed in IP wash/dilution buffer, 3×5 min at 4°C. Input and IP samples were eluted and denatured by boiling in 2x Laemmli buffer/bME for 10 min at 95°C.

Co-IPs were also performed using Flag M2-bound magnetic agarose beads (Sigma) and GFP-Trap beads (ChromoTek). For Flag pull-downs, AID-Usp9x ES cell were used as controls for nonspecific binding to the Flag beads. For GFP pull-downs, the same amount of lysate was added to negative beads (ChromoTek) to control for nonspecific binding to beads. Cells were collected as above but diluted into GFP-Trap dilution buffer (10 mM Tris-HCl pH 7.5, 150 mM NaCl, 0.5 mM EDTA), immunoprecipitated by rotating for 1.5h at 4°C, and washed by 3x fast washes in GFP-Trap dilution buffer. Input and IP samples were denatured as above.

### HA-Ubiquitin Immunoprecipitations

HA-tagged ubiquitin (a gift of the F. Sicheri lab) was overexpressed in ES cells by transfection with Lipofectamine 2000 (Thermo Fisher Scientific), 500 ng per ∼8×10^6^ cells in a 10 cm dish. Water diluted in Lipofectamine was used for mock transfections. Medium was changed the next morning and cells were harvested after 24 hours. Adherent cells were washed twice and then scraped into cold PBS. The resulting cell pellets were weighed and resuspended in 4x volume of RIPA buffer (150 mM NaCl, 1% NP-40, 0.5% Na deoxycholate, 0.1% SDS, 50 mM Tris pH 8) to lyse for 15 min on ice. Pellets were centrifuged at max speed for 10 min at 4°C to remove insoluble material. 100 µl of supernatant was taken for IP and diluted to 500 µl in non-denaturing lysis buffer (20 mM Tris pH 8, 137 mM NaCl, 1% Triton X-100, 2 mM EDTA) plus 2.5 µg of anti-HA antibody (Abcam ab9110). IPs were incubated overnight at 4°C with end-over-end rotation. The next day, immune complexes were bound to 25 µl Protein A Dynabeads (Thermo Fisher Scientific) for 2h at 4°C. Complexes were washed on beads for 3×10 min in IP wash buffer (150 mM NaCl, 10 mM Tris pH 8, 0.5% Na deoxycholate, 1% Triton X-100, 1 mM EDTA, 1 mM EGTA) and eluted in 2x Laemmli buffer/10% β-mercaptoethanol for 10 min at 95°C followed by cold shock on ice. Input samples were collected and denatured in Laemmli buffer to 1x. Samples were removed from beads for western blotting.

For Usp9x catalytic domain expressions, transfections were performed as above but with the addition of 2.5 µg of plasmid (wild-type or C1566S pEF1a-Usp9x_CD-mCherry) and in medium without Pen/Strep. Transfection was checked by mCherry fluorescence the next morning. IPs were performed as above but with the following antibodies instead of HA: Ezh2 at 1:300 (CST #5246), Suz12 at 1:50 (CST #3737), or rabbit IgG at 1:50 (Millipore CS200581).

### Subcellular fractionation

Subcellular fractionation was performed as previously reported^113^. Cell pellets were resuspended in buffer A (10 mM HEPES pH 7.9, 10 mM KCl, 1.5 mM MgCl_2_, 0.34M sucrose, 10% glycerol, 0.1% Triton X-100, 1 mM DTT, and PIs: NaF, PMSF, 1x Halt Protease inhibitor cocktail), incubated 5 min on ice, and centrifuged for 5 min at 1300 *g* at 4°C. The supernatant was taken as the cytoplasmic extract and clarified by centrifugation. Nuclear pellets were washed in buffer A and resuspended in buffer B (3 mM EDTA, 0.2 mM EGTA, 1 mM DTT, and PIs). After 5 min on ice, chromatin pellets were centrifuged for 5 min at 1700 *g*, 4°C. The supernatant was collected as the soluble nucleoplasmic fraction. Insoluble pellets were resuspended in 1x Laemmli buffer containing 5% β-mercaptoethanol and sonicated on a Bioruptor: high power, 30s on, 30s off, 5 min total (Diagenode).

### Western blot analysis

Denatured samples were separated on 4-15% Mini-Protean TGX SDS-PAGE gels (Bio-rad). Protein was transferred to methanol-activated PVDF membranes (Bio-rad) by wet transfer (1x Pierce Transfer Buffer, 10% methanol) or using high molecular weight transfer conditions for the Bio-rad TransBlot Turbo (Bio-rad). Membranes were blocked in 5% milk/TBS-T and incubated with indicated primary antibodies for 1.5h at room temperature or overnight at 4°C. Membranes were then washed and incubated with HRP-conjugated anti-mouse/rabbit secondary antibodies (Jackson Labs) for 1h at room temperature. Proteins were detected by ECL (Pierce) or Clarity (Bio-rad) detection reagents and exposure to X-ray film (Pierce).

For analysis of cell cycle, FUCCI reporter ES cells^33^ were collected by trypsinization and sorted on a FACS AriaII (BD Biosciences) into mCherry+ (G0/G1) and BFP+ (S/G2/M) cell fractions. Cells were pelleted and lysed in RIPA buffer and clarified by centrifugation for 10 min, 13,000 *g* at 4°C. Lysates from the same number of cells were used for western blotting.

### H3K27me3 ChIP-seq

Two biological replicates, consisting of independent clones of AID-Usp9x collected on consecutive days, were collected. 10^6^ cells were sorted and cross-linked in 1% formaldehyde/PBS, rotating for 10 min at room temperature. Cross-links were quenched with glycine (125 mM final) for 5 min at room temperature. Cells were washed 2x in cold PBS, snap frozen, and stored at −80°C. All subsequent steps were performed on ice or at 4°C. Fixed cell pellets were thawed and lysed in 1% SDS, 10 mM EDTA, 50 mM Tris-HCl pH 8 with protease inhibitors (1x Halt Protease inhibitor cocktail, 1 mM PMSF, 1 mM NaF) for 30 min. Chromatin was sheared to 200-500 bp fragments on a Covaris S220 with settings PIP 105, duty 2, cpb 200 for 9 min. Shearing efficiency was confirmed by 1% agarose gel electrophoresis. Chromatin lysates were clarified by centrifugation and diluted 1:10 in dilution buffer (1% Triton X-100, 2 mM EDTA, 167 mM NaCl, 20 mM Tris-HCl pH 8) with protease inhibitors. Inputs were collected at the same time. IPs were performed overnight using 2.5 µg of antibody (CST #9733 H3K27me3 or Abcam ab46540 rabbit IgG) per equivalent of 500,000 cells, rotating at 4°C. The next day, immunocomplexes were precipitated by incubation with pre-washed Protein A Dynabeads (Invitrogen) for 2h. Beads were washed 4×10 min in low-salt buffer (0.1% SDS, 1% Triton X-100, 2 mM EDTA, 150 mM NaCl, 20 mM Tris-HCl pH 8), 1×10 min in high-salt buffer (0.1% SDS, 1% Triton X-100, 2mM EDTA, 500 mM NaCl, 20 mM Tris-HCl pH 8), 1×10 min in LiCl buffer (0.25 M LiCl, 0.5% NP-40, 0.5% Na deoxycholate, 1 mM EDTA, 10 mM Tris-HCl pH 8), and 1x fast in TE. ChIP and input samples were eluted in fresh ChIP elution buffer (1% SDS, 50 mM NaHCO3, 50 mM Tris-HCl pH 8, 1 mM EDTA) and treated with RNase A for 1h at 37°C. Cross-links were reversed by shaking overnight at 65°C with Proteinase K.

Genomic DNA was cleaned up using Qiagen MinElute Reaction Cleanup Kit (Qiagen) and quantified by Qubit (Thermo Fisher Scientific). ChIP efficiency was confirmed by H3K27me3 enrichment relative to IgG IP in qPCR at diagnostic regions. The same amount of chromatin from HEK293 cells was spiked in to equivalent volumes of ChIP eluates (62 pg of spike-in chromatin per 25 µl of ChIP), yielding final concentrations between ∼1-5%. Libraries were constructed from 2.5 ng of DNA and prepared using the NEBNext Ultra II DNA Library Prep Kit for Illumina with 9 PCR cycles (NEB #E7645S, New England Biolabs). Library quality was assessed by High Sensitivity DNA Assay on an Agilent 2100 Bioanalyzer (Agilent Technologies). Samples were sequenced on a HiSeq 4000 using single-end 50 bp reads.

### H3K27me3 ChIP-seq data analysis

Sequencing reads that passed quality control were trimmed of adaptors using Trim Galore! v0.4.0 and aligned to mm10 and hg19 using bowtie2 v2.2.5^114^ with no multimapping. SAM files were converted to BAM files, sorted, and indexed. Normalization factors (NFs) for each sample were calculated as a fraction of input reads^8^. Bam files were deduplicated using picard v2.18.14 MarkDuplicates (http://broadinstitute.github.io/picard). Raw H3K27me3 ChIP-seq data were downloaded as fastq files from NCBI GEO for the indicated datasets. For paired-end samples, only one read was kept per fragment and all samples were trimmed, aligned to mm10, sorted, and deduplicated as above. Deduplicated bam files were analyzed using deepTools v3.3.0 on the command line^115^. Library sizes for normalization were calculated from bam files before deduplication (samtools view -c).

#### Broad peak calling

Deduplicated bam files were converted to scaled bedgraphs using deepTools bamCoverage (options --scaleFactor <NF> --binSize 10 --blackListFileName ENCODE_mm10_blacklist.bed) and then to bed files: awk ‘{print $1” \t” $2” \t” $3” \tid-” NR” \t” $4” \t.”}’. These scaled bed files were used to call broad peaks compared to input using epic2 on the command line (options -gn mm10, -d chrM). Bedtools merge was used to merge peaks within 3kb, and bedtools intersect was used to determine a set of common peaks between replicates. Bam files were converted to scaled bigWigs using deepTools bamCoverage (options --binSize 100 --scaleFactor <NF>). Correlation between replicates was checked by multiBigWigSummary bins and plotCorrelation, and then scaled bw files were merged (bigwigCompare add) for heatmaps. computeMatrix was used to generate coverage of scaled bigwig files over no-auxin peaks (options scale-regions -m 500 --upstream 10000 --downstream 10000 --binSize 100 --missingDataAsZero --skipZeros --sortRegions descend --sortUsing mean --sortUsingSamples 1 -p max). Heatmaps were produced using deepTools plotHeatmap.

TSS profile plots were generated from the output of deeptools plotProfile (--outFileNameData), which was imported into R, processed to average replicates, and then plotted with ggplot2. For H3K27me3 coverage over *Nodal*, bigwig files were downloaded from NCBI GEO^58^. Sample tracks were visualized in Integrated Genome Viewer using bigwig files (IGV v2.3.92).

multiBamSummary was used to count reads falling into non-overlapping 10kb genes across the genome, and read counts were then imported into R. Embryo counts were normalized by library sizes (number of mapped reads in deduplicated bam files), and ES cell data were normalized by spike-in factors. For cumulative distribution plots, reads were counted in non-overlapping 10kb genomic bins using deeptools multiBamSummary (options --smartLabels --blackListFileName --outRawCounts --minMappingQuality 10 -p max). The resulting counts table was imported into R, filtered to remove regions without coverage, scaled with the NFs calculated above, and then plotted using ggplot2 (stat_ecdf). P-values represent Kolmogorov-Smirnov test results using the averages of biological replicates. Counts per bin were adjusted for biological batch (embryo vs. ES cell origin) using ComBat/sva in R^116^ and analyzed by PCA.

For analysis of repetitive elements, H3K27me3 was counted over repetitive elements annotated in the mouse genome (obtained from UCSC RepeatMasker) using featureCounts (options -f -O -s 0 -T 8). In R, we filtered out elements with low coverage, scaled using the NFs calculated above, and calculated the average of replicates. Plots show regions with > 5 normalized counts for hyper-H3K27me3 and < 3 for hypo-H3K27me3 elements, with log_2_(Usp9x-high/no-auxin) > |0.7| as the threshold for enrichment.

## SUPPLEMENTARY FIGURES

**Supplementary Figure 1.**
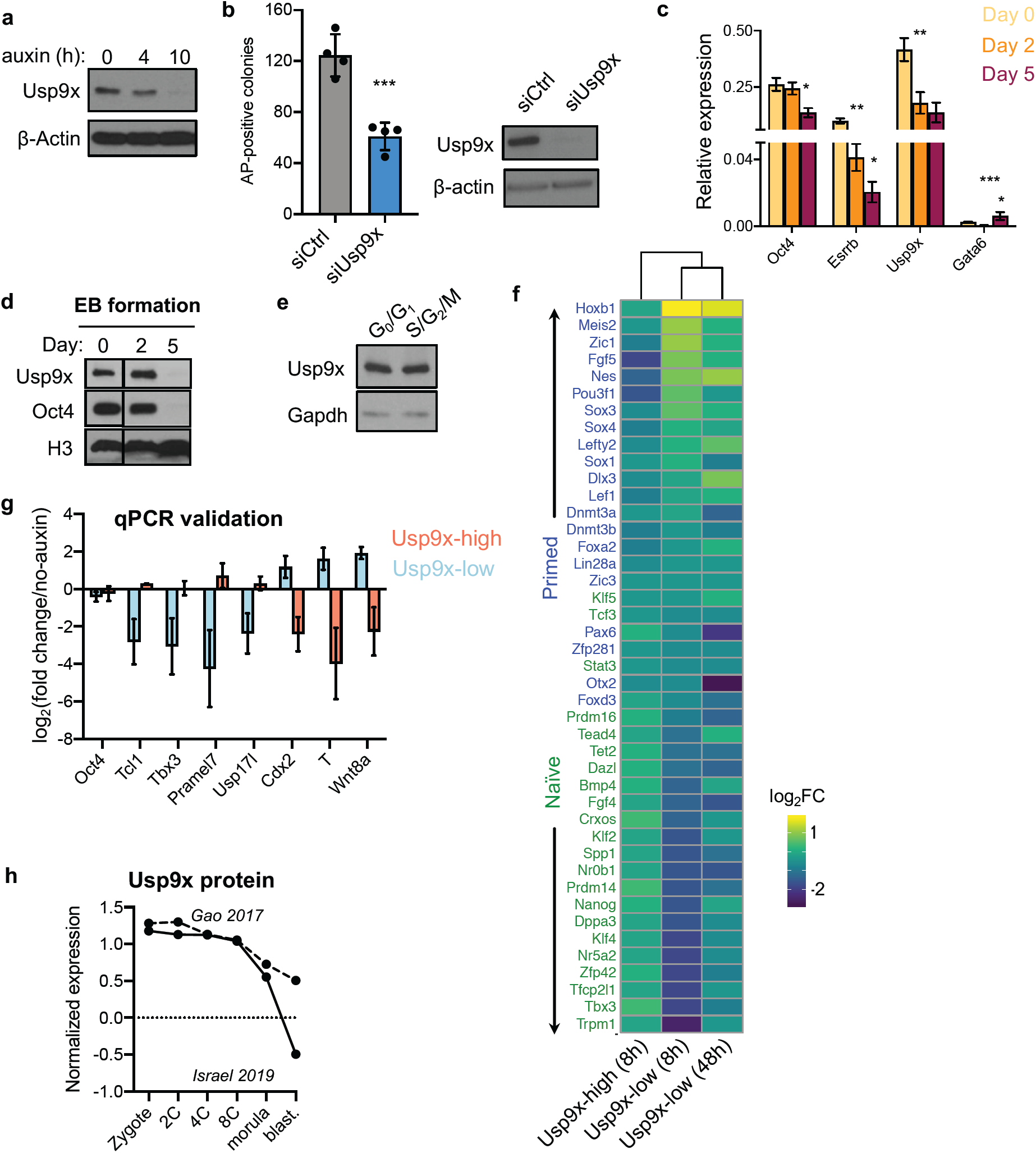
Characterization of Usp9x expression in targeted ES cells and early embryos. **a)** Auxin treatment induces acute depletion of endogenous Usp9x protein over a time course of auxin, 0-10h. **b)** Colony formation assay in control (siCtrl) or Usp9x-depleted (siUsp9x) ES cells with western blot confirming Usp9x knockdown. **c)** Usp9x mRNA expression declines during lineage commitment of ES cells in Embryoid Body (EB) formation. **d)** Usp9x protein expression declines during the initial stages of lineage commitment. **e)** Usp9x expression is comparable between stages of the cell cycle, isolated using a FUCCI live cell cycle reporter^33^. **f)** Relative expression of representative naïve and primed pluripotency genes in the indicated cell states^4,35^. Data are plotted as log_2_ fold-change (FC) in expression relative to controls. **g)** qRT-PCR validation of representative genes from RNA-seq at 8h after auxin. **h)** Usp9x protein expression declines over pre-implantation development, in parallel with the decline in Usp9x mRNA in early development (Fig. 1e). Normalized data are plotted from quantitative proteomic analyses of wild-type embryos^117,118^. Data are mean ± s.d. of 4 replicates and are representative of 3 independent experiments (b), mean ± SD of 3 replicates (c), mean ± s.e.m. of 2 biological replicates (g). Western blots are representative of 2-3 biological replicates. \**P* < 0.05, ** *P* < 0.01, \*\*\**P* < 0.001 by Student’s t-test (b), multiple t tests with Holm-Sidak correction (c).

**Supplementary Figure 2.**
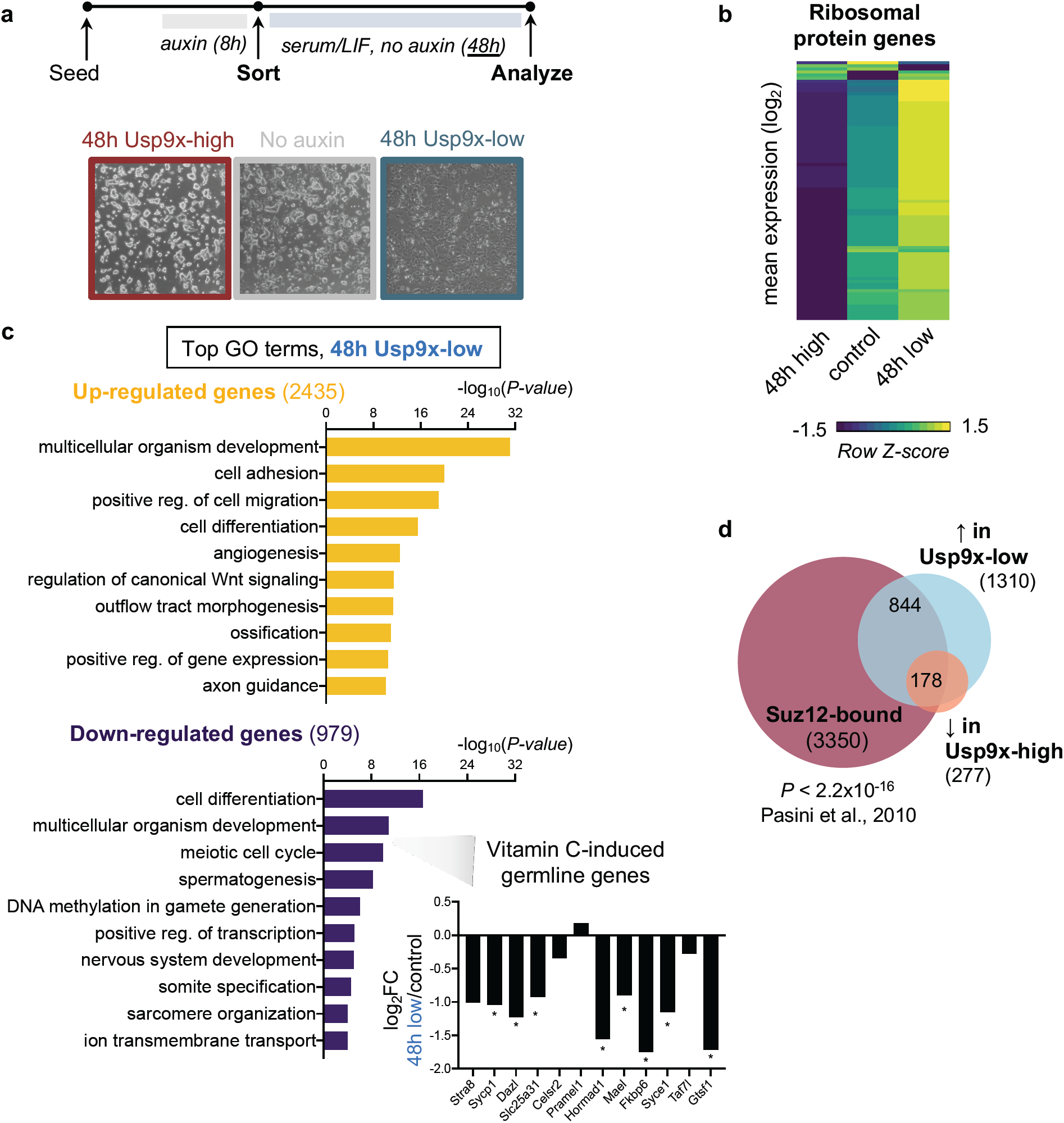
Transcriptional analysis of Usp9x-high and Usp9x-low ES cells at 48h. **a)** Diagram of experiments assessing the ability of sorted Usp9x-high or Usp9x-low ES cells to recover after acute auxin treatment. After recovery, 48h Usp9x-high ES cells form compact colonies and Usp9x-low ES cells adopt heterogeneous, differentiated morphologies. **b)** Relative expression of ribosomal protein genes in Usp9x-high, Usp9x-low, or control cells at 48h. **c)** Gene Ontology (GO) analysis of genes significantly upregulated or downregulated in Usp9x-low ES cells after 48h. Upregulated genes are enriched for differentiation- and development-related GO terms. Downregulated genes are enriched for meiosis- and germline-related GO terms, reminiscent of the hypomethylated state of naïve pluripotency driven by vitamin C addition to 2i ES cell culture. Inset: log_2_ fold-change (FC) in expression of several vitamin C-induced germline genes^2,119^. **d)** Overlap of Suz12-bound genes^43^ with genes DE in Usp9x-high and Usp9x-low ES cells (see Fig. 1f). 178 of 248 overlapping DE genes (72%) are Suz12 targets. Boxplot hinges (b) show the first and third quartiles, with median center line. Data in (c) are the average of 3 replicates. \**FDR* < 0.05 or *P* value as indicated. *P*-values by Wilcoxon rank-sum test (b), output of DESeq2 (c inset), and Fisher exact test (d).

**Supplementary Figure 3.**
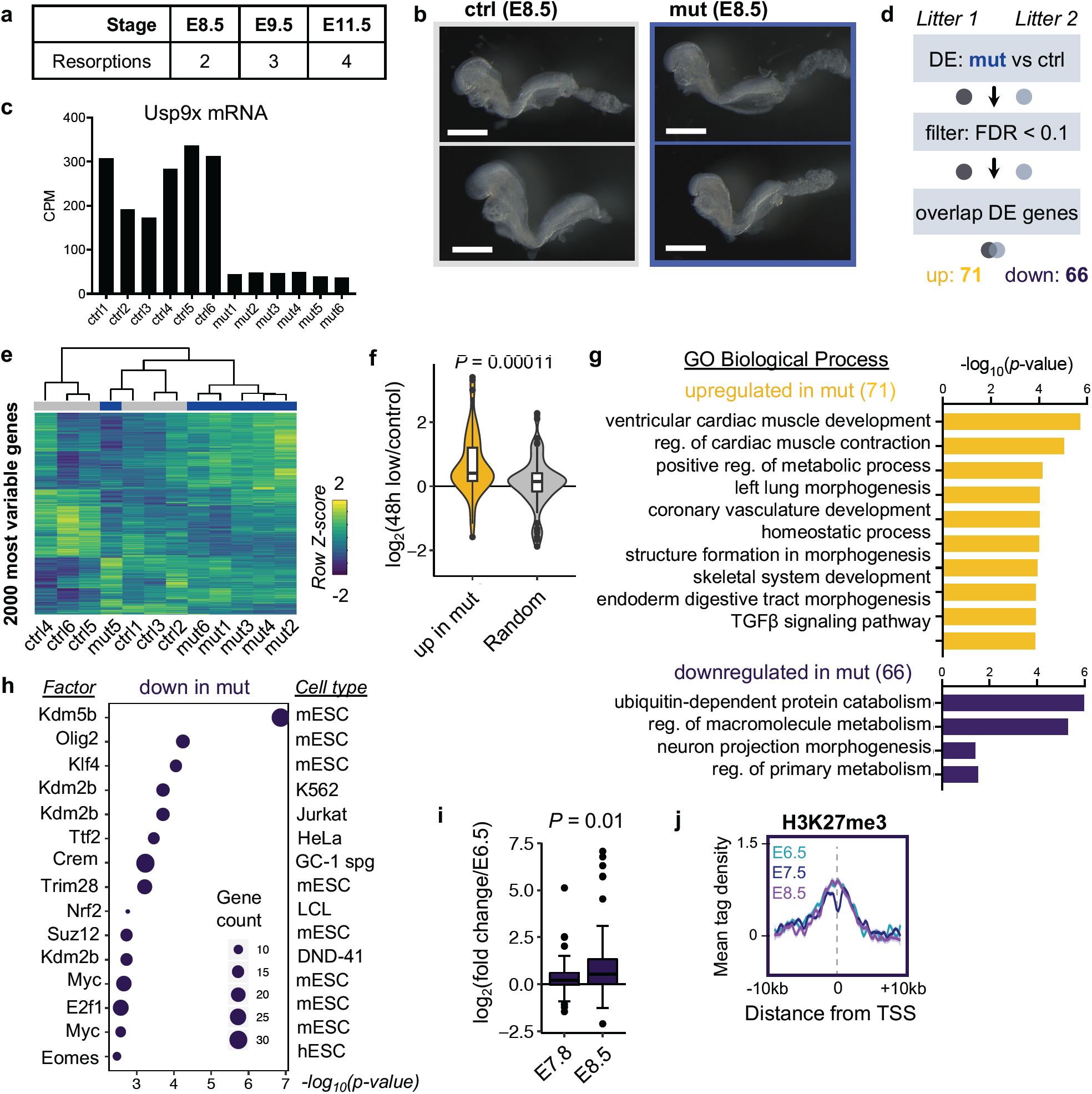
Transcriptional analyses of *Usp9x*-mutant embryos at E8.5. **a)** Number of resorptions counted at the indicated stages (no embryonic material detected in deciduum). **b)** Representative control and mutant embryos at E8.5 used for RNA-seq. Mutants are morphologically indistinguishable from controls at this stage. Scale bar = 500 µm. **c)** Normalized counts confirming low Usp9x mRNA expression in the 6 mutant embryos used for RNA-seq. *n* = 12 embryos from 2 litters were sequenced. **d)** Approach to DE analysis of E8.5 *Usp9x*-mutant transcriptomes. **e)** Unsupervised hierarchical clustering of the top 2000 most variable genes across samples by RNA-seq. Mutants largely cluster away from controls, except for mut5 (litter 2), which clusters with the controls from litter 1. **f)** Boxplot showing that the genes upregulated in *Usp9x* mutants are also up in 48h Usp9x-low ES cells relative to controls, compared to a random subset of the same number of genes. **g)** Top-enriched GO terms for up- and down-regulated genes in *Usp9x* mutants. **h)** Enrichr TF analysis of genes downregulated in *Usp9x-*mutants, similar to Fig. 2e. These genes are targets of repressive chromatin factors, e.g. Kdm5b, Kdm2b, Trim28, and Suz12, in the indicated cell types. **i)** The genes downregulated in *Usp9x* mutants tend to be upregulated by E8.5^56^. **j)** Profile of H3K27me3 ChIP-seq signal during wild-type development over the genes downregulated in *Usp9x* mutants^58^. Boxplot hinges (f,i) show the first and third quartiles, with median center line. Data are mean ± s.e.m. of 2-3 replicates per time point (j). *P* values by Wilcoxon rank-sum test (f, i).

**Supplementary Figure 4.**
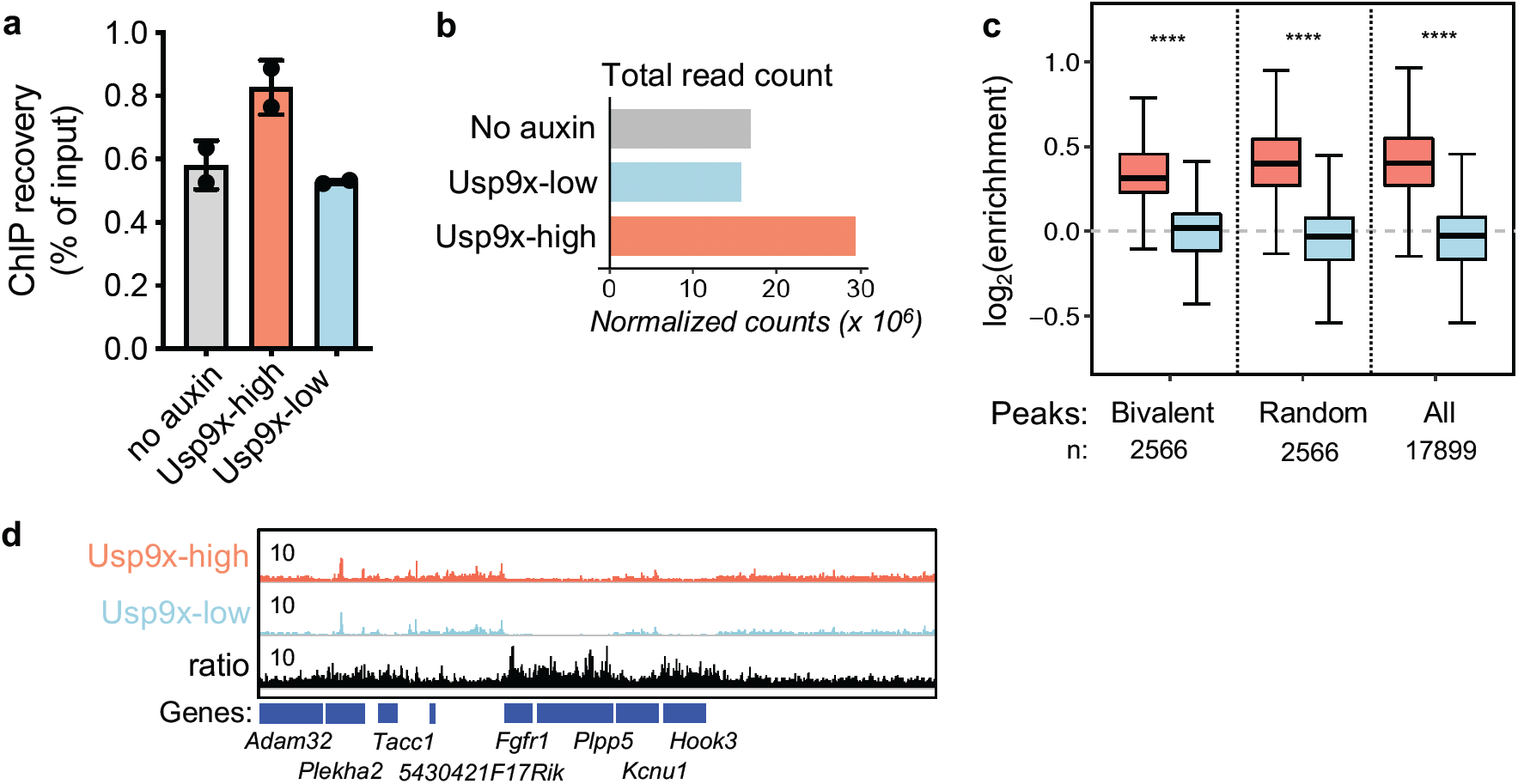
Analysis of H3K27me3 patterns in Usp9x-high and Usp9x-low ES cells. **a)** ChIP recovery before sequencing recapitulates the global gain of H3K27me3 in Usp9x-high ES cells (Fig. 3a). **b)** Spike in-normalized sequencing coverage of H3K27me3 in the indicated cells. **c)** H3K27me3 coverage in Usp9x-high or Usp9x-low cells over bivalent peaks, a random peak set of the same size (2566), or all peaks found in ES cells at baseline (no auxin). **d)** Representative genome browser view of H3K27me3 signal in Usp9x-high and Usp9x-low cells. Elevated H3K27me3 signal in Usp9x-high cells is often observed outside of promoters. Data are mean ± s.d. of 2 replicates (a) or sum of replicates (b-d). Boxplot hinges (c) show the first and third quartiles, with median center line. \*\*\*\**P* < 0.0001 by Wilcoxon rank-sum test (c).

**Supplementary Figure 5.**
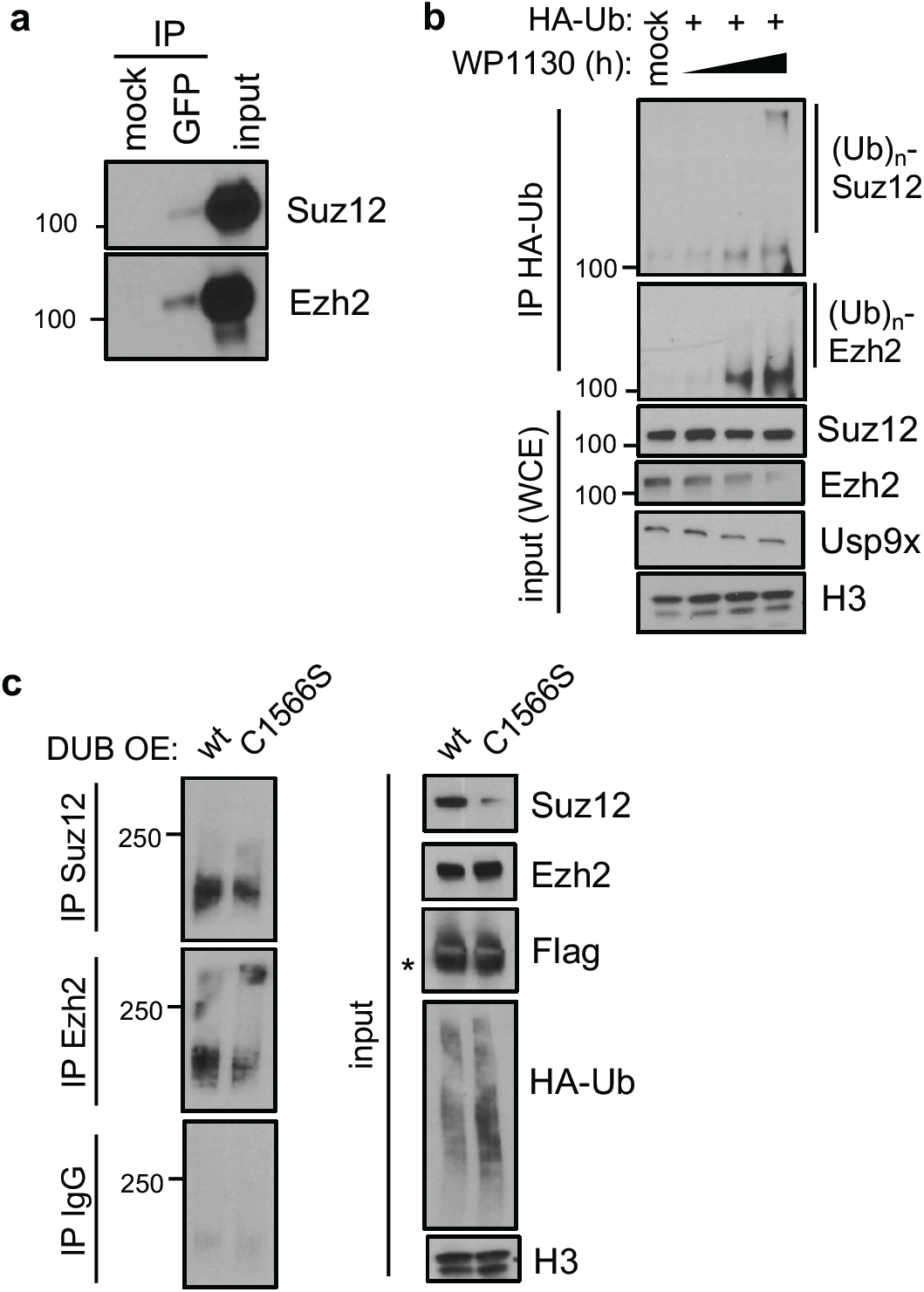
Validation of the Usp9x-PRC2 regulatory interaction. **a)** Co-IP showing that GFP-tagged Usp9x interacts with Suz12 and Ezh2 in AID-Usp9x cells. **b)** Acute catalytic inhibition of Usp9x with the semi-selective inhibitor WP1130 leads to gain of ubiquitin at PRC2 proteins, similar to Fig. 4c. WP1130 treatment ranges from 0-4h. **c)** Comparison of wt versus catalytic-dead Usp9x catalytic domain overexpression but in wild-type ES cells, similar to Fig. 4d. Asterisk (*) designates the expected band size for the Usp9x catalytic domain construct. All western blots are representative of 2-3 biological replicates.

